# Effects of accumulated temperature on growth curve of *Gynostemma pentaphyllum* stem and leaf under the conditions of Simulated light under the forest

**DOI:** 10.1101/2021.03.01.433332

**Authors:** Dun chunyao, Wan Songsheng, Li Shuanglong, Zeng yong, Wu Daikun

## Abstract

The effects of accumulated temperature on the growth curve and leaf number growth curve of *Gynostemma pentaphyllum* were studied. The growth curve of twigs and strands were studied by curve regression analysis. The results showed that the growth of *Gynostemma pentaphyllum* Leaf growth curve was optimized, and the growth curve of stem and leaf of *Gynostemma pentaphyllum* was established, and the curve was fitted and analyzed. Three - dimensional model of the effect of accumulated temperature on the stem length and leaf number of *Gynostemma pentaphyllum*. The results show that the stem length L and the accumulated temperature t of the strands are in the logistic function, the mathematical model was L=-112.69/(1+(t/892.1)^3.31^)+130.54, the number of gibbere leaf number n is logistic function with the accumulated temperature t, The model was n=-27.86/(1+(t/1159.77)^0.26^)+30.37, and the growth model could reflect the dynamic growth of *Gynostemma pentaphyllum* growth. The experimental results provide a theoretical basis for choosing suitable habitat for *Gynostemma pentaphyllum* under the forest.

## INTRODUCTION

*Gynostemma pentaphyllum* (Thunb.) Makino) is a perennial herbaceous vine of the genus *Gynostemma* in the *Cucurbitaceae* family. Distributed in the Qinba Mountains and Wuling Mountains in China, traditional Chinese medicine and minority areas in China are often used as anti-inflammatory drugs to treat colds, coughs, etc. *Gynostemma* flavonoids and saponins have anti-cancer effects, and *Gynostemma* polysaccharides can eliminate inflammation and antioxidant The activity(Tsai,Y.C.Lin,C.L., et al, 2010). *Gynostemma pentaphyllum* has formed a large industry in Shaanxi, Hubei and other province of China. Because of its shade-loving characteristics, it can be planted under forest in mountainous areas and is an economic crop with great potential. The stems and leaves of *Gynostemma pentaphyllum* are mainly used directly. The stems and leaves of *Gynostemma pentaphyllum* grow rapidly. The growth rate can be directly expressed by measuring the elongation of the stem and the number of leaves to establish a growth curve model.

The accumulated temperature required for specific plants in the growth stage is relatively stable. There are many studies on predicting the growth and development period of plants based on the physiological development time, but currently they are mainly focused on the research of field crops, and there are few reports on medicinal plants at home and abroad (PENG JIANZHONG,1995). Accumulated temperature is an important factor that affects the physiological development time and growth. The physiological development time can be used as an intermediate factor to clarify the functional relationship between accumulated temperature and growth, and then establish a growth curve model during a specific growth period (ZHANG ZHIYOU,2012). The commonly used models for studying plant growth curves include Logistic function and Gompertz function. The ideal growth curve model plays an important role in guiding the cultivation of medicinal plants under the forest. It can not only master the basic law of the growth and development of medicinal plants, but also combine with environmental condition data to analyze the growth status and predict the growth trend (ZHANG YU et al.,2010), it is convenient to choose the under-forest environment suitable for the growth of medicinal plants. At present, no research has been found to track the growth rate and accumulated temperature of *Gynostemma pentaphyllum*.

## MATERIALS AND METHODS

### Test materials

The test material is “Enwuyemi” *Gynostemma pentaphyllum* seedlings, provided by Hubei Xianzhitang Biological Technology Co., Ltd., collected from its base in Lichuan City, Enshi Prefecture, Hubei Province, China. *Gynostemma pentaphyllum* seedlings are selected to maintain the same height, stem thickness, number of leaves, etc., and cultured with perlite as a solid substrate with improved Hoagland medium. During the vegetative growth period (15 March to 21 June), cultivate at room temperature, measure the temperature at 02, 08, 14, and 20 o’clock every day, and calculate the average daily temperature.

### Test method

Observe the growth status of *Gynostemma pentaphyllum,* measure it every 3-9 days, measure the data of the stem length and the number of leaves of *Gynostemma pentaphyllum* in different growth periods, and analyze the functional relationship between the two.

The illumination in the cultivation room is set to 500-1000lux, which simulates the underforest environment, and the humidity and other environments are stable. Record the temperature of the cultivation room at 4 time points, including 2:00, 8:00, 14:00, 20:00 o’clock every day The daily average temperature is obtained, and the time to determine the length of the stem and the number of leaves of *Gynostemma pentaphyllum* is a calculation cycle to eliminate the error influence of individual extreme conditions.

### Statistical analysis

Pre-process the original data with Excel software. SPSS 19 software and Oringe8.0 software were used to analyze the function relationship between accumulated temperature and *Gynostemma pentaphyllum* stem length and leaf number data, establish the stem length and temperature change curve, leaf number and temperature change curve, and calculate the accumulated temperature and stem elongation and leaf by time correlation The functional relationship between the numbers, establish the growth and development model of *Gynostemma pentaphyllum.*

## Results and analysis

### The relationship between Gynostemma pentaphyllum stem length and accumulated temperature

*Gynostemma pentaphyllum* stem elongation and growth are positively correlated with accumulated temperature. According to the curve estimation, the trend of Logistic model fits well with the relationship between *Gynostemma pentaphyllum* stem length (L) and accumulated temperature (t), as shown in Fig. 1, and its calculation formula is shown as Equation 1 after fitting. The fit R^2^ was 0.9893. Excluding the rate fluctuation caused by the factors of light intensity and sunshine hours, with the accumulation of accumulated temperature, its growth rate first increases and then decreases. When the environmental accumulated temperature is about 867.5 °C, its growth rate is the fastest.

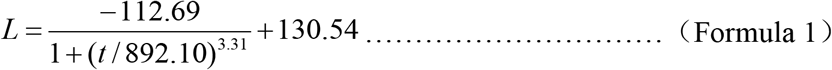

**Figure 1.**
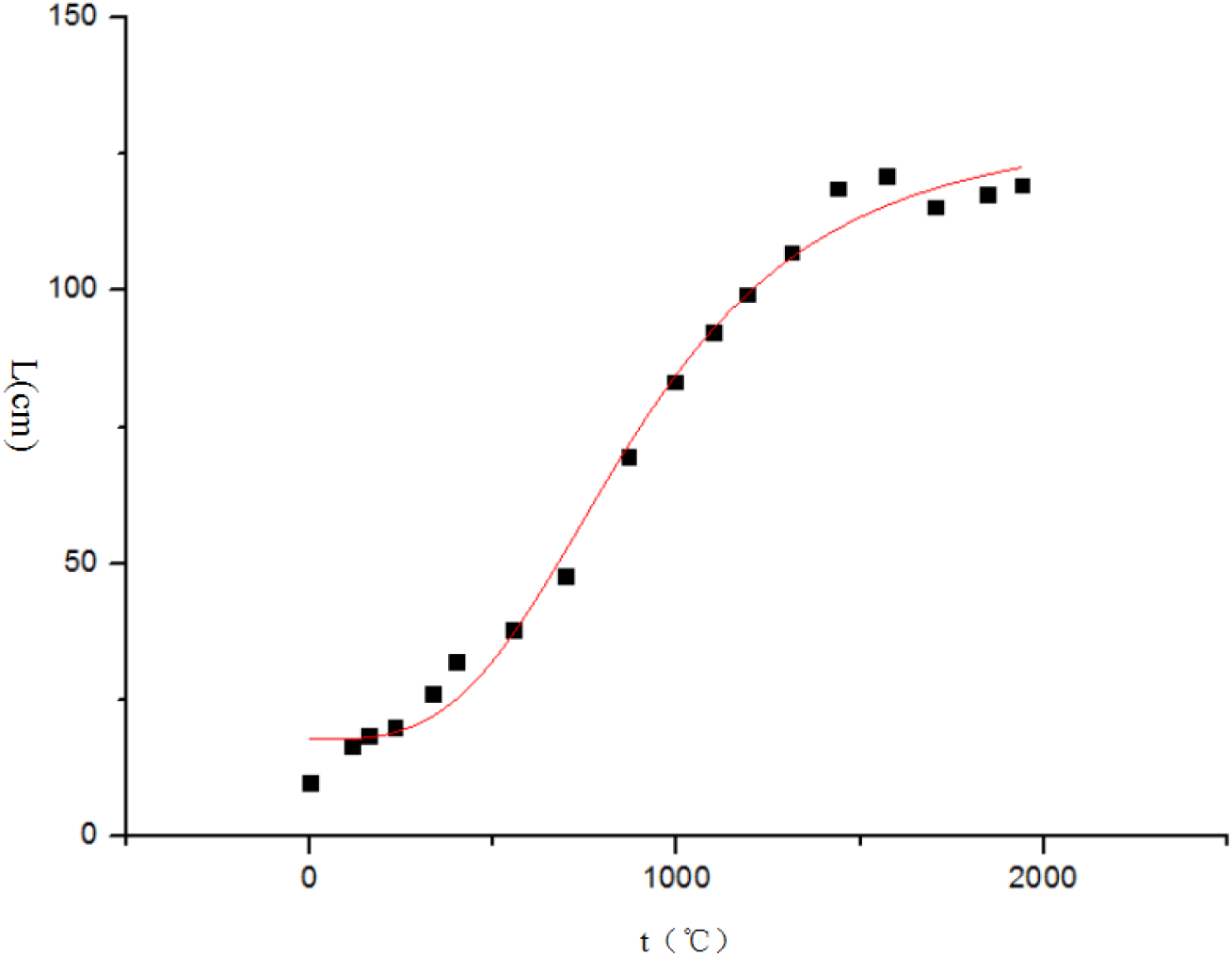
Gynostemma pentaphyllum stem length and accumulated temperature

### The relationship between the number of Gynostemma pentaphyllum leaves and the accumulated temperature

The number of *Gynostemma pentaphyllum* leaves is positively correlated with the environmental accumulated temperature. According to the curve estimation, the trend of the Logistic model fits well with the relationship between *Gynostemma pentaphyllum* stem length (n) and accumulated temperature (t). After fitting, the calculation formula was shown in Equation 2, and the degree of fit R^2^ was 0.9919, as shown in Figure 2. With the accumulation of accumulated temperature, the growth rate of the number of leaves increased first and then decreased. When the accumulated temperature of the environment was around 994.5°C, its growth rate was the fastest, but the influence of accumulated temperature on the growth rate of the number of leaves was less than that of the stem growth.

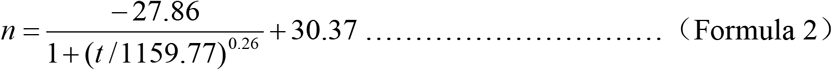

**Figure 2.**
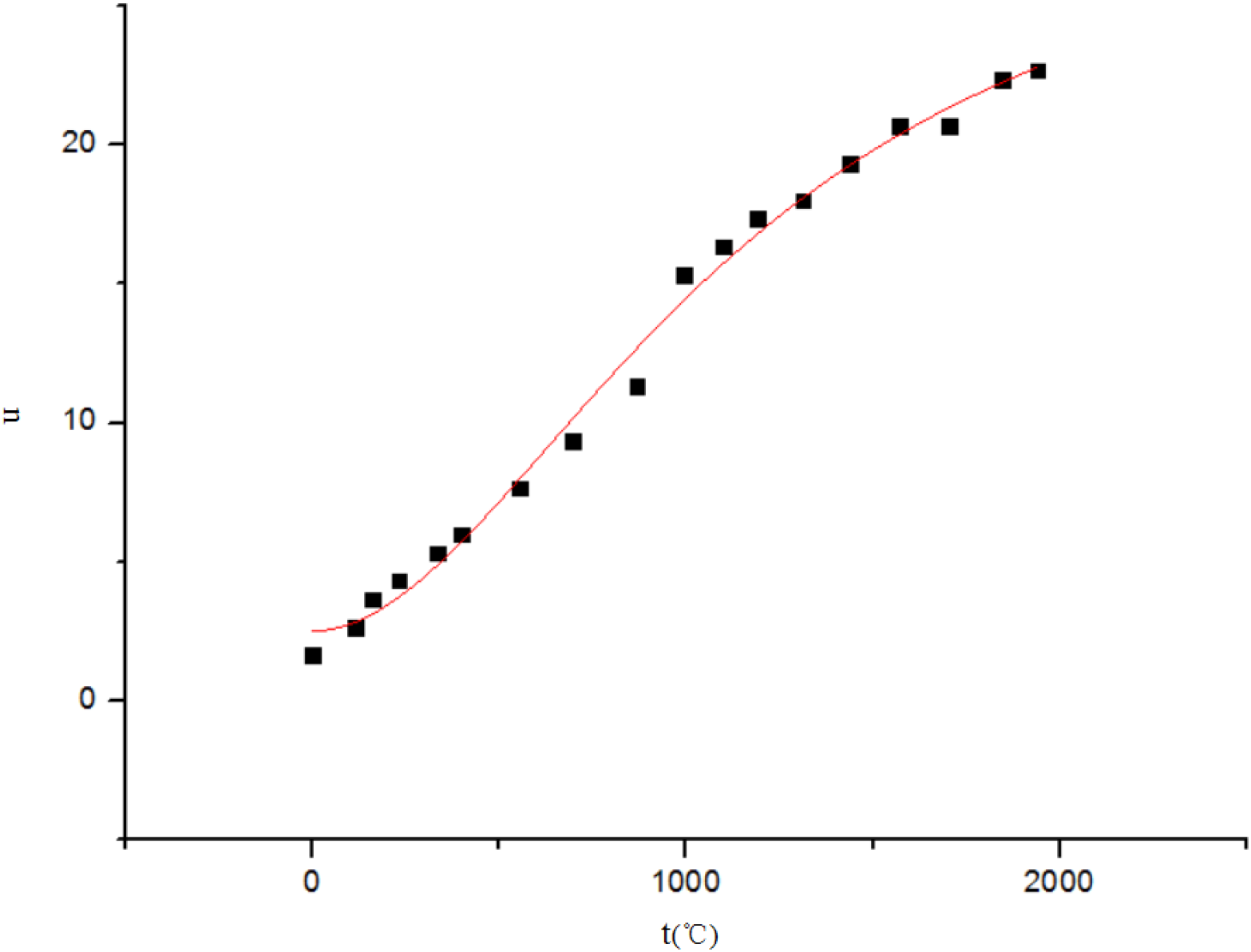
Gynostemma pentaphyllum leaf number and accumulated temperature

### Construction of three-dimensional model of leaf number, stem length and accumulated temperature

In order to more intuitively describe the direct relationship between the number of leaves, stem length and accumulated temperature, a three-dimensional model is established with the accumulated temperature as the x-axis, the stem length as the y-axis, and the number of leaves as the z-axis, as shown in Figure 3., The fit R^2^ was 0.9933. With the increase of accumulated temperature, the stem length and the number of leaves of *Gynostemma pentaphyllum* increased, and the number of leaves showed a positive correlation with the stem length.

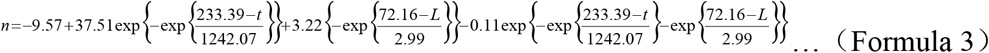

**Figure 3.**
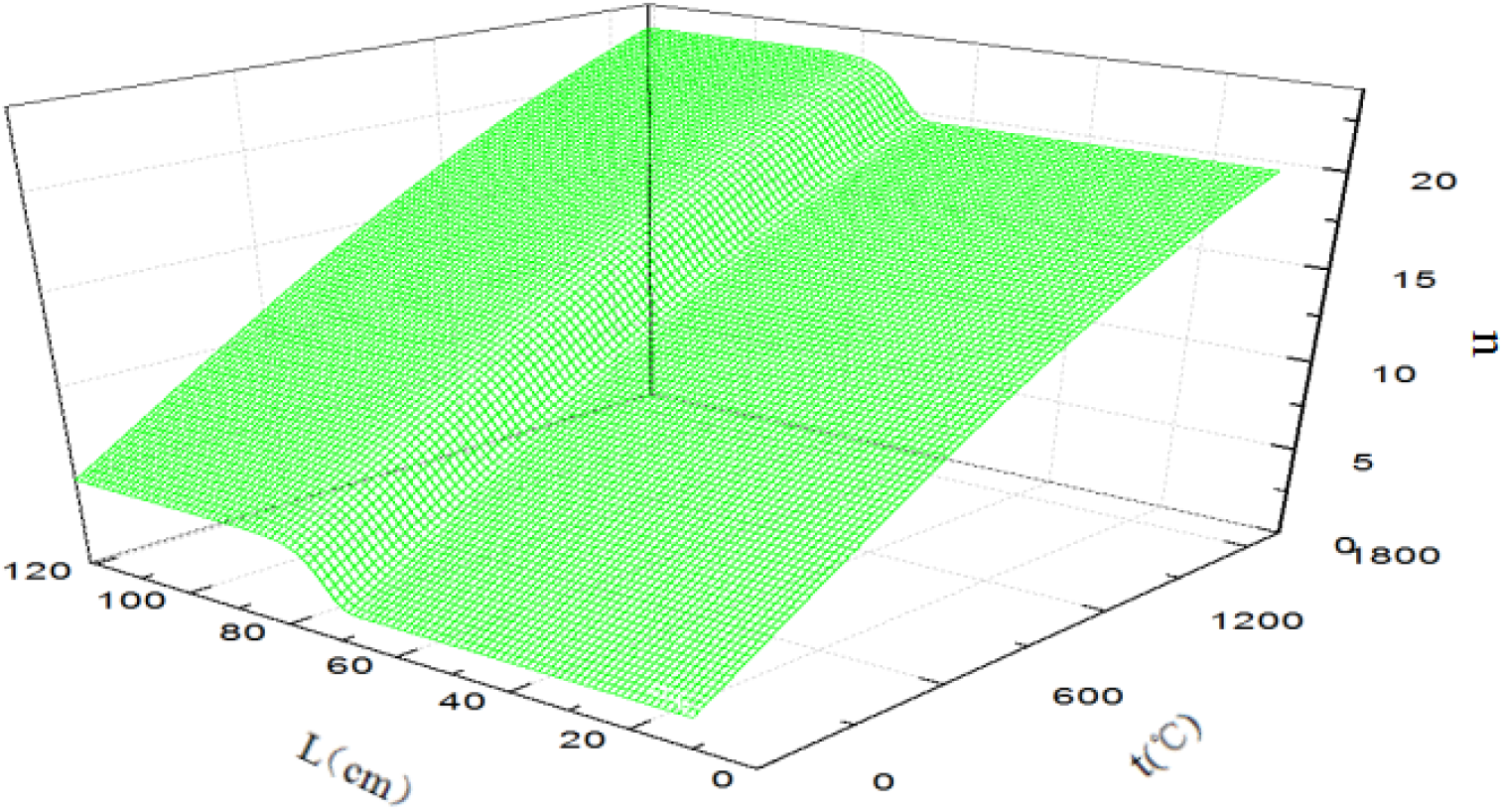
Gynostemma pentaphyllum stem length and accumulated temperature

## DISCUSSION AND CONCLUSION

Establishing the growth curve of *Gynostemma pentaphyllum* stem and leaf-accumulated temperature can guide *Gynostemma pentaphyllum* planting and production. Regression analysis was performed on growth time and accumulated temperature. The results are shown in Figure 4. The R^2^ was 0.995 after inspection. The growth time and accumulated temperature are linearly related, and accumulated temperature and growth time show a high degree of correlation. Therefore, it is feasible to use accumulated temperature instead of growth time to fit the growth curve and simulate the growth of *Gynostemma pentaphyllum.* In actual forestry production, under different altitude zones, different latitude zones and different microclimate conditions, there are great differences in the growth environment of understory plants. Existing studies only use the growth time as a reference system to simulate plant growth conditions. Therefore, exploring the relationship between environmental temperature and the growth of *Gynostemma pentaphyllum* stems and leaves can provide a better reference for choosing the environment for cultivation under the *Gynostemma pentaphyllum* forest.

**Figure 4.**
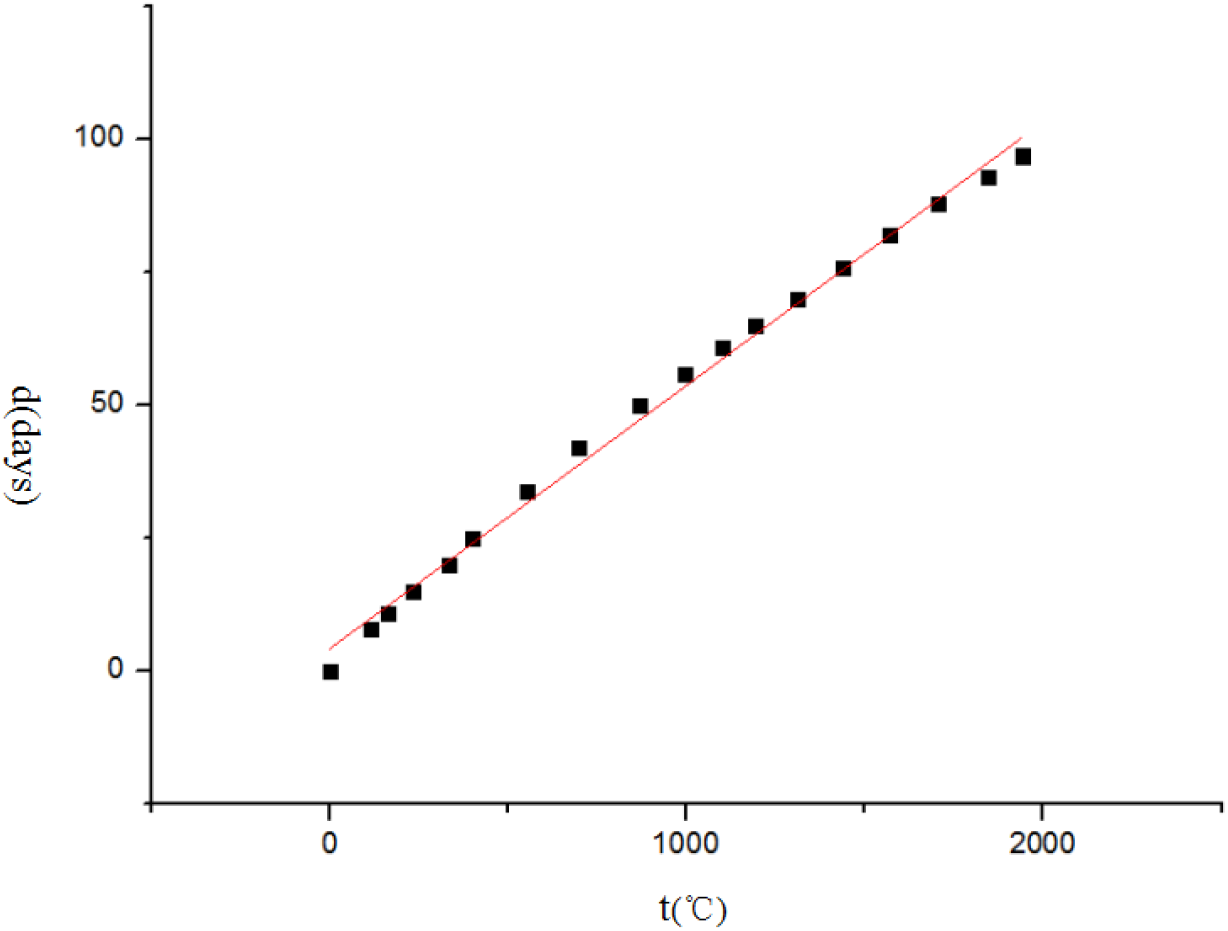
Accumulated temperature and growth days

In production, the first tender leaves of *Gynostemma pentaphyllum* can be picked in June each year. Products made from this material have higher economic added value. Therefore, this article only simulates the growth of *Gynostemma pentaphyllum* during the vigorous growth period from March to June. The growth of *Gynostemma pentaphyllum* in the north of China is relatively large from June to August (WANG GUANGLU & Li BANGQIN,1994), so the relationship between the growth of stems and leaves of *Gynostemma pentaphyllum* and the accumulated temperature in large-scale production in different regions needs further study. *Gynostemma pentaphyllum* is mainly used in leaves, and its main biologically active ingredient is saponins. Therefore, the content of saponins directly affects the quality of *Gynostemma pentaphyllum* products. At present, there are few studies on the relationship between environmental temperature and Gypenoside accumulation rules, and relevant research should be strengthened. The substrate used in this study is perlite and Hoagland nutrient solution is used for cultivation. The soil conditions are very different from the understory environment. Further research is needed to establish the growth model of *Gynostemma pentaphyllum* under forest conditions.

The growth model fitted in this paper can better reflect the dynamic changes of *Gynostemma pentaphyllum* during the vigorous growth period, and establish a three-dimensional model of the effect of accumulated temperature on the stem length and leaf number of *Gynostemma pentaphyllum*. The experimental results provide a theoretical basis for large-scale cultivation of *Gynostemma pentaphyllum* under forest.

## ACKNOWLEDGEMENTS

The paper is supported by the science and technology project D20190020 of Enshi Autonomous Prefecture. We thank Hubei University for Nationalities for providing the basic conditions for related experiments and Professor Zheng Xiaojiang for his guidance.

## DECLARATION OF CONFLICT OF INTEREST

We have no conflict of interest to declare.

## AUTHORS’ CONTRIBUTIONS

All authors contributed equally for the conception and writing of the manuscript. All authors critically revised the manuscript and approved of the final version.

